# LIPID ANCHORING OF ARCHAEOSORTASE SUBSTRATES AND MID-CELL GROWTH IN HALOARCHAEA

**DOI:** 10.1101/863746

**Authors:** Mohd Farid Abdul-Halim, Stefan Schulze, Anthony DiLucido, Friedhelm Pfeiffer, Alexandre W. Bisson Filho, Mechthild Pohlschroder

## Abstract

The archaeal cytoplasmic membrane provides an anchor for many surface proteins. Recently, a novel membrane anchoring mechanism involving a peptidase, archaeosortase A (ArtA) and C-terminal lipid attachment of surface proteins was identified in the model archaeon *Haloferax volcanii*. ArtA is required for optimal cell growth and morphogenesis, and the S-layer glycoprotein (SLG), the sole component of the *H. volcanii* cell wall, is one of the targets for this anchoring mechanism. However, how exactly ArtA function and regulation control cell growth and mor-phogenesis is still elusive. Here, we report that archaeal homologs to the bacterial phosphatidylserine synthase (PssA) and phosphatidylserine decarboxylase (PssD) are involved in ArtA-dependent protein maturation. *H. volcanii* strains lacking either HvPssA or HvPssD exhibited motility, growth and morphological phenotypes similar to those of ∆*artA*. Moreover, we showed the loss of covalent lipid attachment to SLG in the ∆*hvpssA* mutant and that proteolytic cleavage of the ArtA substrate HVO_0405 was blocked in the ∆*hvpssA* and ∆*hvpssD* strains. Strikingly, ArtA, HvPssA, and HvPssD GFP-fusions co-localized to the mid position of *H. volcanii* cells, strongly supporting that they are involved in the same pathway. Finally, we have shown that the SLG is also recruited to the mid cell prior to being secreted and lipid-anchored at the cell outer surface. Collectively, our data suggest haloarchaea use the mid cell as the main surface processing hotspot for cell elongation, division and shape determination.

## Introduction

Microbial cell surface proteins play critical roles in many important biological processes, including bioenergetics, mediation of intercellular communication, nutrient uptake, surface adhesion, and motility. Cell surface proteins also play important roles in cell elongation and shape maintenance, but how this is achieved in archaea is not well understood (1).

The structural organization of cellular surfaces is one important readout of how cells coordinate growth, morphogenesis, and division. In both bacteria and eukaryotes, a multitude of growth modes have been characterized, with cells inserting new envelope material almost all along the cell surface (2), bipolarly (3), unipolarly (4), and in some cases, different modes can be interchangeable (5, 6). In the case of archaea, which lack a peptidoglycan cell wall, glycosylated S-layer and other proteins are commonly the sole component of the cell envelope (7, 8), where they typically show a 2D crystal-like arrangement. This poses an interesting problem for archaeal surface protein organization, and currently there is no data about the mechanisms of archaeal cell elongation control (9).

While many proteins are anchored to the cell surface via transmembrane (TM) domain insertion into the membrane, some are anchored through covalent N-terminal attachment of a lipid moiety (8). Recently, a novel mechanism was discovered whereby proteins are anchored to the membrane through a lipid moiety covalently attached to a processed C-terminus. In archaeal cells, processing and lipid modification of these C-terminal anchored proteins are mediated by enzymes known as archaeosortases, with archaeosortase A (ArtA) of the model archaeon *Haloferax volcanii* being the most studied example (10–13). Proteins recognized and processed by *H. volcanii* ArtA contain a distinct C-terminal tripartite structure consisting of a conserved PGF motif, followed by a hydrophobic domain and then a stretch of positively charged residues. Molecular biological and biochemical analyses determined that ArtA does indeed process a diverse set of proteins, including both Tat and Sec substrates, that have been shown to play roles in motility and mating (10, 12). Most notably, this includes the S-layer glycoprotein (SLG), which is the sole component of the *H. volcanii* cell wall.

A previous *in silico* study by Haft and coworkers noted that in *Methanosarcina acetivorans* C2A, *Methanosarcina mazei* Gö1, as well as several other methanogens, the *artA* gene is located next to the gene that encodes an archaeal homolog of bacterial phosphatidylserine synthase (PssA) (14). Based on the degree of sequence similarity and its substrate specificity, the archaeal PssA homolog belongs to the PssA subclass II, similar to PssA found in gram-positive bacteria such as *Bacillus subtilis*, as opposed to PssA of gram-negative bacteria, such as *Escherichia coli*, which belong to subclass I (15).

Work in *B. subtilis* has elucidated most of the biochemistry involved in the reaction catalyzed by PssA, which involves the transfer of a diacylglycerol moiety from a CDP-phosphatidyl lipid to L-serine to make phosphatidylserine (16, 17). Phosphatidylserine can subsequently be decarboxylated to phosphatidylethanolamine by the enzyme phosphatidylserine decarboxylase (PssD), which has been characterized from *Sinorhizobium meliloti* and *B. subtilis* (17, 18). However, unlike bacterial PssA, *in vitro* study of the archaeal PssA homolog from *Methanothermobacter thermautotrophicus* (MTH_1027) revealed that this protein catalyzes the transfer of the archaetidic acid moiety of CDP-archaeol onto the hydroxyl group of L-serine to form the polar lipid archaetidylserine (CDP-2,3-di-O-geranylgeranyl-sn-glycerol:L-serine O-archaetidyltransferase) (15).

Mirroring the phosphatidylethanolamine biosynthesis reaction in bacteria, it was postulated that archaetidylserine could also undergo decarboxylation to archaetidylethanolamine by an archaeal PssD homolog, a putative archaetidylserine decarboxylase.

Distant homologs to PssA and PssD are encoded in the *H. volcanii* genome, which we refer to as HvPssA (HVO_1143) and HvPssD (HVO_0146). In this study, we show that HvPssA and HvPssD are involved in ArtA-dependent C-terminal protein maturation, which involves proteolytic cleavage and lipid anchoring. An interplay between ArtA, HvPssA and HvPssD is further supported by their colocalization at mid cell. These analyses reveal, to the best of our knowledge for the first time, that cell elongation happens from mid cell in archaea.

## Results

### Synteny of *artA*, *pssA* and *pssD* genes in *Methanosarcina strains*

Based on the juxtaposition of the genes encoding the archaeosortase (*artA*) and the putative membrane lipid biosynthesis archaetidylserine synthase (*pssA*) in *Methanosarcina* (14), we hypothesized that the *H. volcanii* homologs of PssA (HvPssA) might be involved in lipidation of ArtA substrates.

Upon further *in silico* analyses, we were intrigued that in several *Methanosarcina* species the *artA* and *pssA* genomic region flanked a homolog of the *pssD* gene, probably encoding archaetidylserine decarboxylase, suggesting they could be involved in consecutive steps of a proposed archaetidylethanolamine biosynthetic pathway. However, in *H. volcanii*, the genes encoding ArtA (HVO_0915), HvPssA (HVO_1143) and HvPssD (HVO_0146) are not clustered in the same genomic neighborhood (Fig. 1).

**Figure 1:**
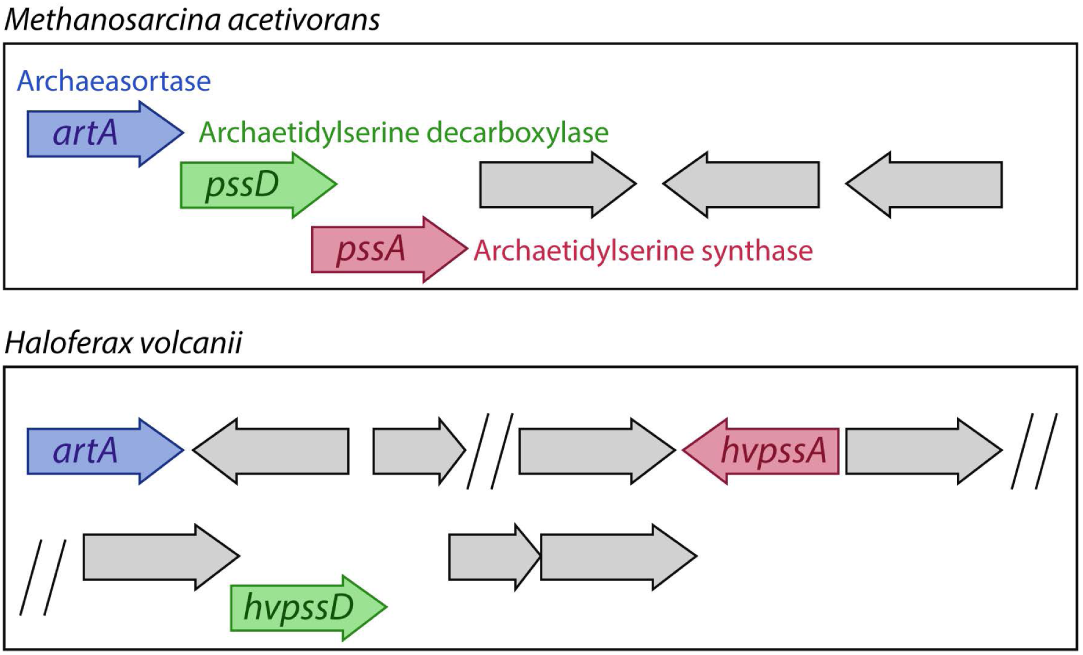
Schematic representation of *artA*, *pssA*, and *pssD* distribution across Euryarchaeota. **(top)** *M. acetivorans* and **(bottom)** *H. volcanii* genomic organization of *artA*, and genes encoding homologs to PssA (HvPssA) and PssD (HvPssD).

It should be noted that while the *Methanosarcina spp.* and *H. volcanii* PssA and PssD homologs share a significant similarity, these proteins have only 30-35% sequence identity to the *in vitro* characterized enzymes from*. M. thermoautotrophicus*, *S. meliloti*, and *B. subtilis* (15, 17, 18). Thus, it is possible that the *H. volcanii* proteins may act on variants of the substrates processed by these experimentally characterized homologs.

### *H. volcanii* Δ*hvpssA* and Δ*hvpssD* cells exhibit growth, morphology, and motility phenotypes similar to those of the Δ*artA* strain

In order to determine whether HvPssA and/or HvPssD are involved in the archaeosortase-dependent processing pathway, we generated *H. volcanii hvpssA* and *hvpssD* deletion mutants (Fig. S1), using the pop-in pop-out method (19).

We had previously shown that the *H. volcanii* Δ*artA* strain exhibits various severe phenotypic defects (e.g. poor growth, atypical morphology, impaired motility), perhaps due, at least in part, to defective processing of the SLG, an ArtA substrate (10). The Δ*hvpssA* and Δ*hvpssD* strains exhibit a growth defect similar to that of the Δ*artA* strain, as compared to the H53 parent strain used in these studies (Fig. 2A). For all genes, normal growth is rescued by complementation, expressing the deleted gene in trans from a plasmid. Moreover, the Δ*hvpssA* and Δ*hvpssD* strains are partially impaired in motility, the defect, however, being less severe than in Δ*artA*. While no halo is observed after 5 days of Δ*artA* incubation at 45°C, a small halo is formed by the Δ*hvpssA* and Δ*hvpssD* strains (Fig. 2B).

**Figure 2:**
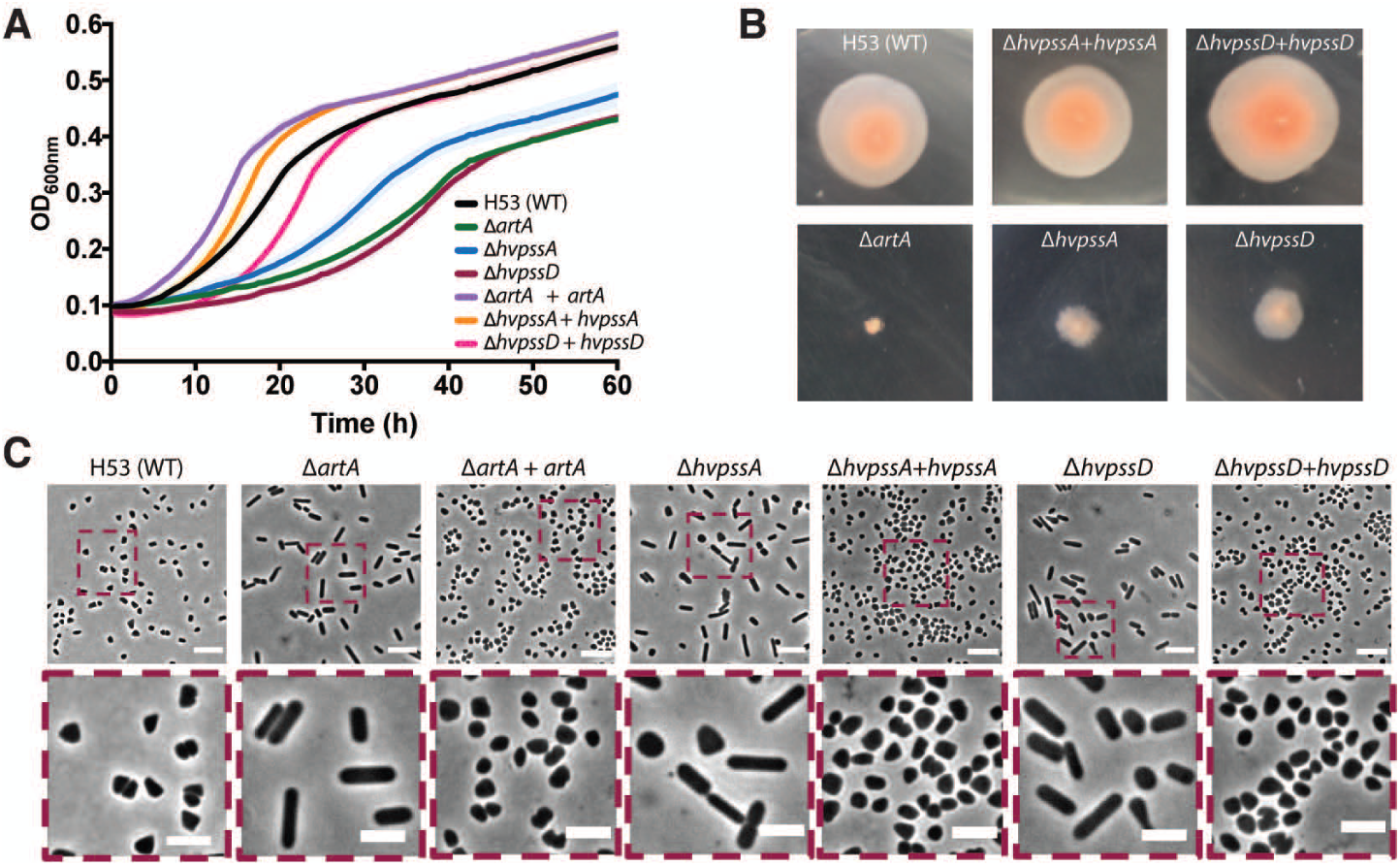
Absence of HvPssA or HvPssD leads to defects in growth, cell morphology, and motil-ity. **(A)** Growth curves: H53 (WT), Δ*artA*, Δ*hvpssA*, and Δ*hvpssD* cells were grown with shaking in 96-well plates with a total volume of 200 µl of liquid semi-defined CA medium and growth of six biological replicates was monitored at OD_600_ with recordings every 30 min. For com-plementation analysis, *artA, hvpssA* or *hvpssD* were expressed from pTA963, under the trypto-phan-inducible P*tnaA* promoter. Both the H53 and deletion strains were transformed with an empty pTA963 plasmid as a control. **(B)** Motility assay: H53, *artA*, *hvpssA*, or *hvpssD* deletion and complementation strains from individual col-ony on solid agar plates were individually stab-inoculated with a toothpick into semi-solid 0.35% agar in CA medium supplemented with trypto-phan, followed by incubation at 45 ^o^C. **(C)** Phase-contrast images were taken from cells during mid-exponential growth phase (OD_600_ 0.3). Scale bars represent 5 µm.

Light microscopic examination of the parental strain at mid-exponential growth stages shows predominantly disk-shaped cells, while the Δ*artA* cells exhibit a predominantly rod-shaped phenotype (Fig. 2C). Again, the Δ*hvpssA* and Δ*hvpssD* strains show a similar but less severe phenotype. The vast majority of cells from cultures are rods, but we have observed a few disk-shaped cells in liquid cultures for each of these strains. This phenotype is fully complemented and disk-shaped cells are observed when HvPssA or HvPssD are expressed in trans in the Δ*hvpssA* and Δ*hvpssD* strains, respectively (Fig. 2C). Thus, Δ*hvpssA* and Δ*hvpssD* strains exhibit phenotypes similar to the Δ*artA* strain for three independent physiological effects, supporting the hypothesis that the encoded proteins are involved in the ArtA-dependent processing pathway.

### HvPssA and HvPssD are required for SLG lipid modification

To confirm that the drastic cell morphology transitions and the phenotypic similarity between the Δ*hvpssA/*Δ*hvpssD/*Δ*artA* mutants are due to the inhibition of the covalent lipid modification of ArtA substrates, we investigated the lipidation of the SLG mediated by ArtA (11). Initially, as an indirect analysis, we examined the effect of *hvpssA* and *hvpssD* deletions on SLG electrophoretic mobility in an LDS-PAGE gel, as a mobility shift is observed in an *artA* deletion strain (11).

While the similarity of Coomassie-stained band intensities for SLG isolated from the Δ*artA*, Δ*hvpssA* and Δ*hvpssD* strains and their parent strains indicate a similar SLG abundance, electrophoretic mobility demonstrates similar migration shifts of the SLG isolated from all three deletion strains as compared to SLG from the parent strain (Fig. 3A). To corroborate this observation, we set out to directly measure lipid modification of SLG in the Δ*hvpssA* strain.

**Figure 3:**
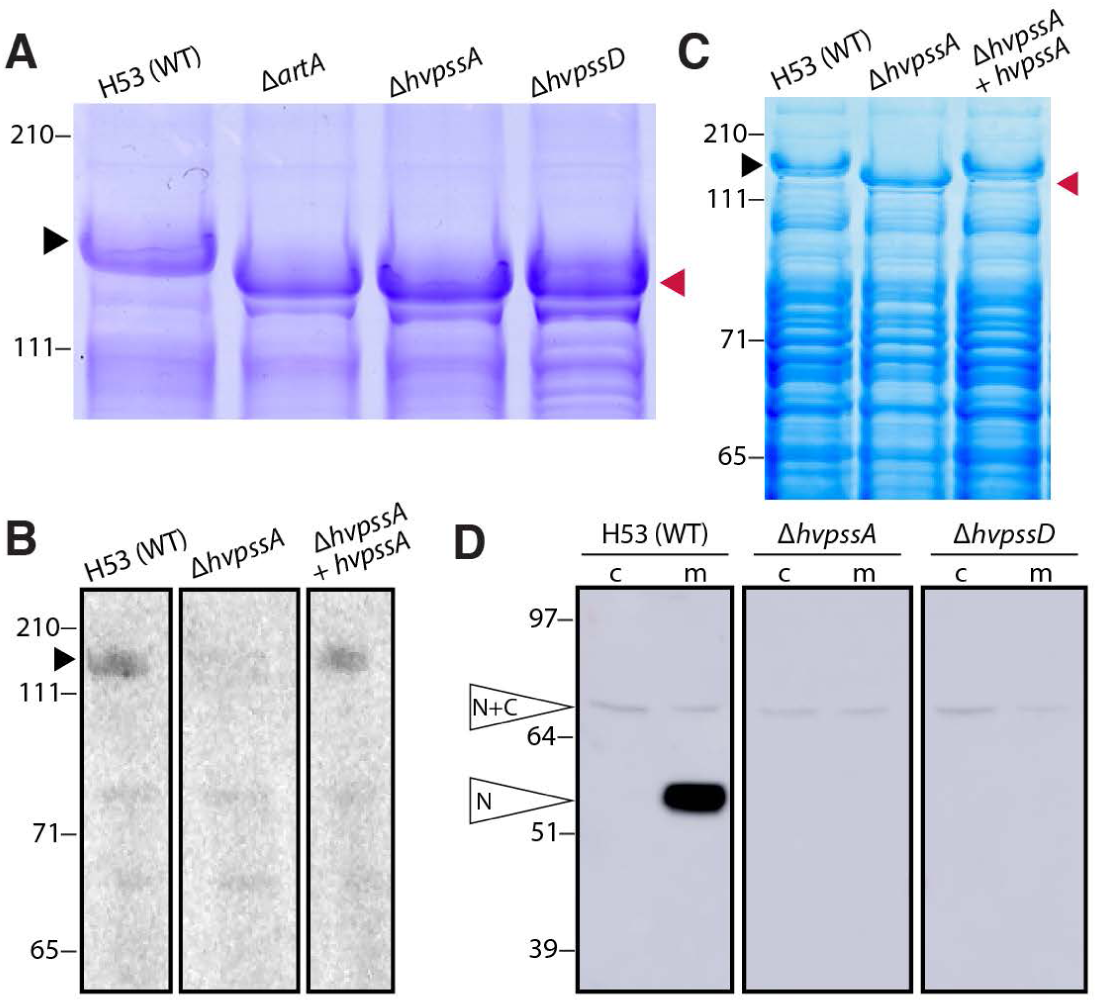
HvPssA and HvPssD are critical for HVO_0405 C-terminal processing and SLG lipidation. **(A)** Coomassie-stained LDS-PAGE gel of cell extracts from *H. volcanii* H53 (wt), Δ*artA*, Δ*hvpssA*, and Δ*hvpssD* strains. The Δ*artA*, Δ*hvpssA* and Δ*hvpssD* SLG (red arrow-head) exhibited a mobility shift compared to the wt SLG (black arrow-head). **(B)** Fluorography of protein extracts isolated from H53 (wt), Δ*hvpssA* and *hvpssA* complementation (Δ*hvpssA + hvpssA*) cells grown in the presence of 1 µCi/ml ^14^C mevalonic acid. Significant la-beling of SLG (black arrowhead) is only detected in the WT and *hvpssA* complementation (Δ*hvpssA* + *hvpssA*) extracts. **(C)** Coo-massie-staining of the gel used for fluorography. The SLG mobility shift in Δ*hvpssA* (red arrowhead) is reverted upon *hvpssA* expression *in trans*. **(D)** Western blot analysis of cytoplasmic (c) and membrane (m) fractions of H53 (wt), Δ*hvpssA*, and Δ*hvpssD* strains expressing, *in trans*, HVO_0405-6xHis. The N-terminal domain of HVO_0405 was detected using anti-HVO_0405-N-term antibodies. HVO_0405 not processed by ArtA and N-terminal HVO_0405 protein processed by ArtA are labeled as “N+C” and “N”, respectively. The C-terminal do-main, which carries a His-tag, has not been analyzed in this experiment. Numbers indicate molecular mass in kilodaltons.

These experiments proved that lipid labeling of the SLG with radiolabeled mevalonic acid, an archaeal lipid precursor, is severely impaired in the Δ*hvpssA* strain as compared to the parent strain (Fig. 3B) and that this phenotype can be complemented by expressing HvPssA *in trans*.

This is conclusive evidence for HvPssA being closely or even directly coupled with ArtA-dependent protein lipidation. Having run out of this label (and not being able to easily obtain more), we could not perform the equivalent experiment for the Δ*hvpssD* strain. However, because the phenotypes of the Δ*hvpssD* strain closely match those of the Δ*hvpssA* strain in all other assays, including the indirect gel-based assay for SLG modification (Fig. 3C), it is highly likely that HvPssD is also involved in ArtA-dependent SLG lipidation.

### HvPssA and HvPssD are required for proteolytic ArtA-substrate processing

To further support that HvPssA/HvPssD-dependent lipidation is required for ArtA-dependent C-terminal processing, we investigated the C-terminal proteolytic cleavage of a second ArtA substrate. As a reporter, we used the strain-specific domain fusion protein HVO_0405, with its centrally located cleavage site, because the cleaved and uncleaved versions of HVO_0405 can be easily distinguished by LDS-PAGE separation and subsequent immunostaining (12). In contrast, this ArtA substrate is cleaved in the corresponding parent strain as is evident from our Western blot analysis of membrane fractions using anti-HVO_0405 antibodies (Fig. 3D and Fig. S2).

### Protein-internal serine residues as an alternative substrate for HvPssA

Before we became aware that the gene encoding a PssD homolog is located between *artA* and *pssA* in several *Methanosarcina* strains, we considered the possibility that internal Ser residues of substrates might be targeted by HvPssA. This would reflect a restricted shift of substrate specificity from free serine to a protein-internal Ser residue, a shift feasible at only 30% protein sequence identity. This possibility was experimentally addressed. Ser to Ala replacement mutations in HVO_0405 supported this hypothesis (Fig. S3). However, with the detection of HvPssD as an additional player, this hypothesis became less likely, as this probable decarboxylase requires a free carboxyl group, which is lacking in internal Ser residues. However, the observed lack of substrate cleavage for HVO_0405 S463A could indicate that the processing site might be at or close to the mutated Ser, which is only five residues upstream of the PGF motif.

### Mid-cell localization of ArtA, HvPssA, and HvPssD promote cell site specific lipidation *in vivo*

Given the dependence of ArtA activity on HvPssA and HvPssD, we considered that ArtA activation required the direct or indirect interaction with HvPssA/HvPssD and/or the product of HvPssA/HvPssD catalysis. To test these hypotheses, we investigated the localization of ArtA, HvPssA and HvPssD in *H. volcanii* live cells. Strikingly, ArtA, HvPssA, and HvPssD tagged with msfGFP localize at mid cell (Fig. 4A). As controls, we also imaged a tagged version of FtsZ1, that has previously been shown to localize to mid cell and was speculated to participate in cell division (20), as well as msfGFP not fused to any protein. FtsZ1-msfGFP shows an almost identical localization pattern compared to ArtA, HvPssA and HvPssD, while the msfGFP protein by itself is not recruited to mid cell (Fig. 4A). Furthermore, time-lapses of cells cultivated within micro-fluidics suggest that ArtA/HvPssA/HvPssD proteins are recruited to mid cell right after daughter cells are born, and persist for most of the cell cycle, including during cytokinesis (Fig. 4B, blue arrowhead) (Movie S1).

**Figure 4:**
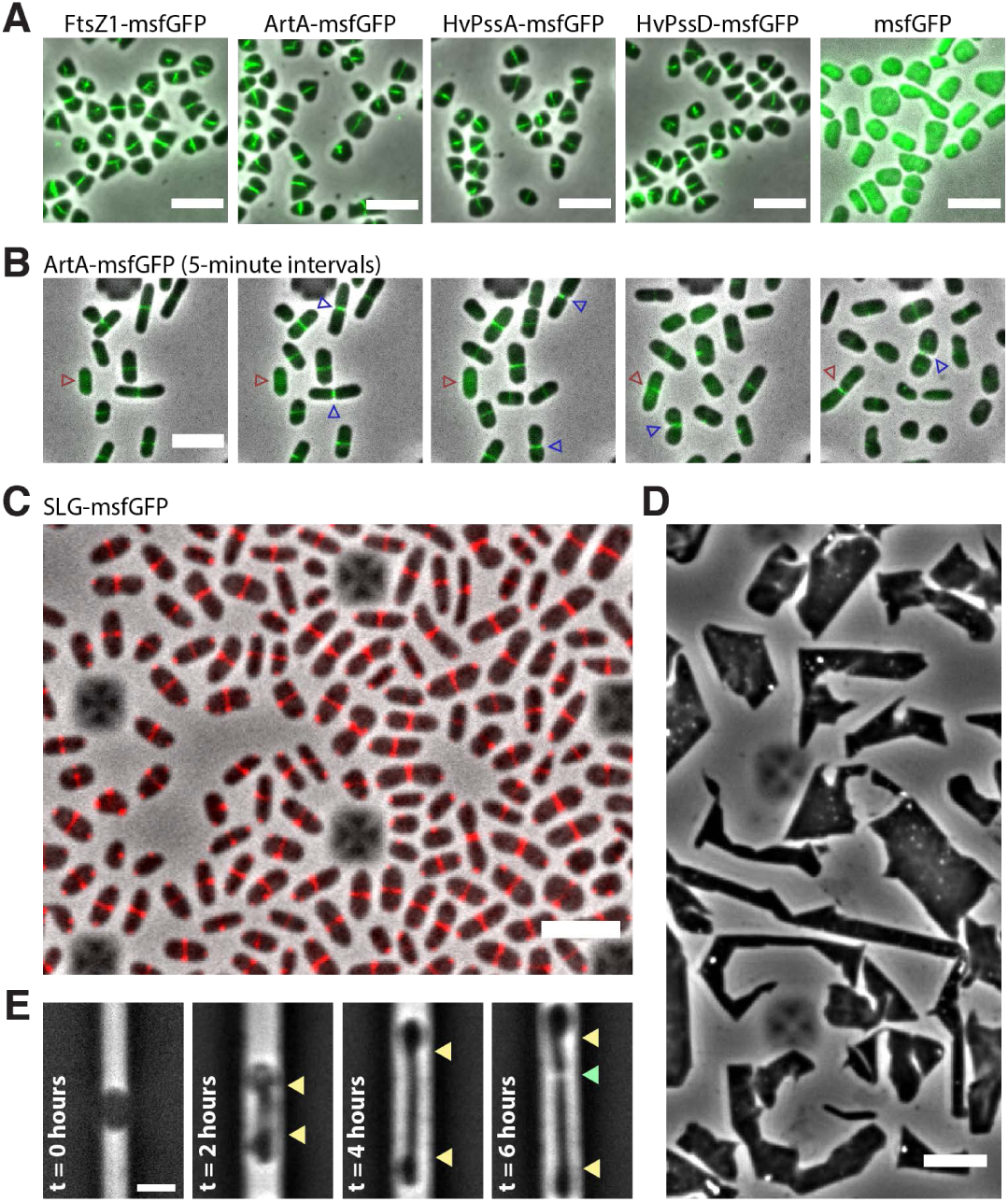
Mid-cell localization of the lipid-anchoring and processing ma-chinery in *H. volcanii*. (A) Snapshots of merged phase contrast (grey) and FITC (green) channels of cells expressing FtsZ1-msfGFP, ArtA-msfGFP, HvPssA-msfGFP, HvPssD-msfGFP and soluble msfGFP. Cells were immobilized under 0.5% agarose pads prepared with CA media. (B) Time lapses of cells growing inside CellAsic microfluidic device. Images of merged phase contrast (grey) and FITC (green) were taken every 5 minutes for 12 hours. Blue arrowheads indicate cell divi-sion events, while red arrowheads label one example of a cell elongat-ing only after the arrival of ArtA-msfGFP to mid cell. (C) Snapshot of SLG-msfGFP (red) mid-cell localization. (D) Phase-contrast images of *H. volcanii* cells under prolonged overexpression of the SLG-msfGFP fusion. (E) *H. volcanii* cells reshape and elongate preferentially at the mid cell during protoplasting recovery. Cells were loaded into the microfluidic chamber, S-layer was chemically removed by addition of 1 mg/mL Proteinase K and recovered with fresh media (t=0 hours). Yel-low arrowheads indicate cell area extended until cell division (green arrowhead). Scale bars represent 5 µm.

Considering that deletions of *artA*, *hvpssA* and *hvpssD* each drastically perturbed growth (Fig. 2A), induced cells to stay in rod-like shape (Fig. 2C), and do not seem to play an essential role in cell division, we hypothesize lipid-anchoring of ArtA substrates specifically at mid cell might be important for cell elongation and morphogenesis. Interestingly, we also noticed a correlation between the presence of ArtA/HvPssA/HvPssD at the mid cell and actual cell elongation in the cell population (Fig. 4B, red arrowhead). To investigate this further, we expressed a second copy of the SLG tagged with msfGFP, a GFP variant shown to be fluorescent upon Sec-dependent transport in bacteria (21). Interestingly, the SLG-msfGFP fusion accumulated at the mid cell site instead of localizing around the cell envelope (Fig. 4C, Movie S2). Additionally, overexpression of SLG-msfGFP fusion caused severe growth and morphological defects (Fig. 4D). These results suggest that the SLG-msfGFP fusion could be not properly secreted but still interact with the Sec system, thus blocking the secretion of untagged SLG and other Sec substrates; alternatively, secreted SLG-msfGFP may be interfering with the 2D proteinaceous crystal array. However, independent of whether this SLG construct was secreted or not, our results strongly suggest that nascent SLG is targeted specifically to the mid cell.

Lastly, we investigated the morphological transitions in *H. volcanii* protoplast cells during *de novo* S-layer synthesis. If the mid-cell confined ArtA/HvPssA/HvPssD are in fact promoting lipidation of recently secreted SLG molecules at the cell surface, then one would be able to observe mid-cell localized reshaping during protoplast recovery. As expected, protoplasts generated by the addition of Proteinase K either within microfluidics or in bulk cultures assume a round-like shape (Fig. 4E, left panel), suggesting the S-layer might be the structure that ultimately determines the archaeal cell shape. As the protease is washed out and replaced by fresh media, cells rapidly reshape exclusively from midcell position (Fig. 4E). Altogether, our observations suggest that the midcell area in *H. volcanii* cells is not only dedicated to cell division, but also a central hub for outbound cell extension and other cellular processes.

## Discussion

Our data confirmed the hypothesized involvement of the lipid biosynthesis enzyme homologs HvPssA and HvPssD in the C-terminal post-translational modifications of ArtA substrates. With respect to physiological effects, ∆*hvpssA* and ∆*hvpssD* showed similar but slightly less severe phenotypes than those resulting from the deletion of *artA* (Fig. 2). We furthermore experimentally determined the effects of *hvpssA* and *hvpssD* deletions on proteolytic cleavage and lipid labeling (Fig. 3).

Since the gene encoding HVO_0405 resulted from the fusion of two previously independent genes, this protein provided us with an excellent tool for the analysis of ArtA-related proteolysis in *H. volcanii*. Not only the mature protein but also the released protein fragment is large and stable enough to be detected by immunostaining (12). This allowed us to clearly demonstrate that ArtA-dependent proteolytic cleavage is blocked when either *hvpssA* or *hvpssD* are deleted in *H. volcanii*. This block could be bypassed by plasmid-based gene complementation. These results strongly suggest that lipidation and proteolysis are intricately connected with proteolytic cleavage only occurring if the modifying lipid and/or HvPssA/HvPssD is present. By sequence homology, *H. volcanii* HvPssA is predicted to be involved in lipid biosynthesis, specifically, the generation of the polar lipid archaetidylserine from CDP-archaeol. However, the functionally characterized homolog (from *M. thermautotrophicus*) is only distantly related (30%-35% sequence identity). We considered that HvPssA might have a related, but distinct, function, with HvPssA acting on a Ser residue within a protein rather than on free serine. Ser to Ala replacement mutations in HVO_0405 (Fig. S3) supported this hypothesis. However, with the confirmed involvement of HvPssD, which requires the availability of a free carboxyl group, this hypothesis became unlikely. Instead, in combination, HvPssA and HvPssD might be generating archaetidylethanolamine.

Interestingly, a recent characterization of a bacterial rhombosortase, a non-homologous analog of archaeosortase (22), showed a direct involvement of glycerophosphoethanolamine-containing moiety in the process. Analogously, lipid-attached ethanolamine may be directly involved in the membrane anchoring of ArtA-substrates. In this scenario, instead of being directly involved in the ArtA-mediated substrate cleavage and/or lipid anchoring, HvPssA and HvPssD catalyze the final steps in a pathway that generates archaetidylethanolamine, a substrate required by this process. This opens the way for another hypothesis regarding the ArtA reaction mechanism. In this scenario, ArtA acts similar to sortase A in bacteria: ArtA cleaves the substrate through thioesterification, forming a thioester acyl‐ enzyme intermediate, which is consistent with the identification of Cys-173 as an active site residue (13). The nucleophilic attack of an amine resolves this intermediate but instead of a pentaglycine branched lipid II, the reactive amine nucleophile is archaetidylethanolamine (Fig. 5A). Such a mechanism would directly result in a covalently modified protein C-terminus. While archaetidylethanolamine lipid was reported to be absent from *Halobacterium salinarum* or *Haloarcula marismortui* (23), it is present in *H. volcanii* and several other haloarchaea albeit with varying abundance (24).

**Figure 5:**
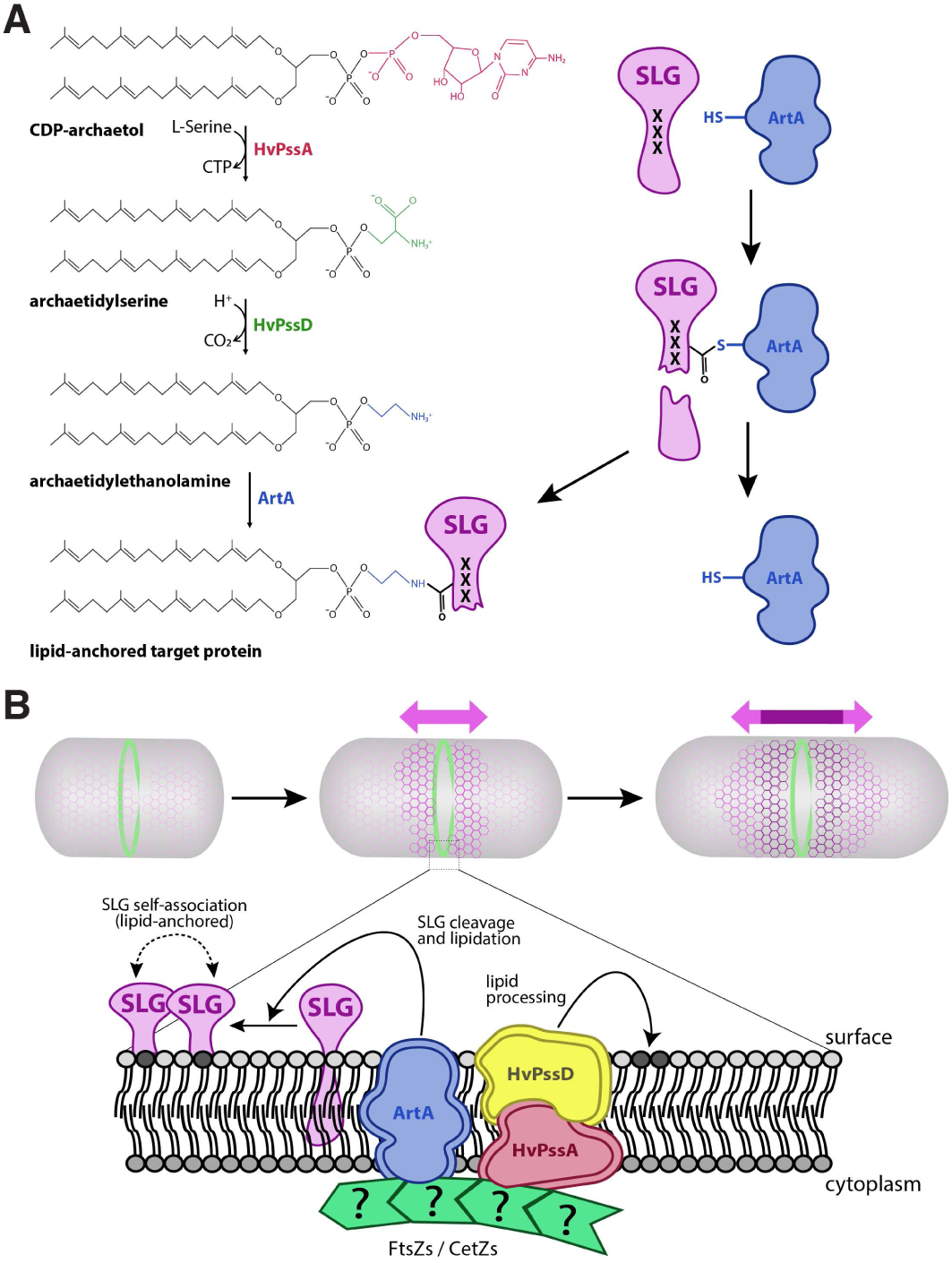
A model for lipid attachment and cell growth involving HvPssA, HvPssD, and ArtA. **(A)** In our speculative model, CDP-ar-chaeol is converted to archaetidylethanolamine in two steps involving HvPssA and HvPssD. ArtA acts as a peptidase and covalently links its active site cysteine to a newly generated C-terminus of its target protein, simultaneously releasing the C-terminal peptide. Then, the free amino group of ethanolamine attacks the thiocarboxylate, which marks the covalent attachment of the target protein to the ArtA active site cysteine. This results in covalent attachment of the lipid to the C-terminus of the target protein as a carboxamide, simultaneously re-leasing ArtA. The process of cleavage and lipidation is dependent on HvPssA and HvPssD, either by binding of archaetidylethanolamine to ArtA or by protein-protein interaction between ArtA and HvPssA/ HvPssD. **(B)** Recruitment of ArtA/HvPssA/HvPssD to mid cell promotes anchoring of surface proteins and insertion of new SLG into the S-layer at mid cell, contributing to cell elongation and division.

As lipid analysis does not cope with protein-bound lipids, the low concentration of archaetidylethanolamine is not surprising, even though the SLG is highly abundant and archaetidylethanolamine may be used as its membrane anchor. Nevertheless, a detectable amount of archaetidylethanolamine in *H. volcanii* membrane suggest the functional roles of HvPssA and HvPssD: catalyzing the synthesis of archaetidylserine and its decarboxylation to archaetidylethanolamine, respectively. These enzymes perhaps associate with or even form a complex with ArtA, resulting in a majority of the synthesized archaetidylethanolamine to be immediately used to modify the SLG and other ArtA substrates for their membrane anchoring upon C-terminal processing. Thus, only a small amount may be left free in the membrane. The lipid attached to an EDTA-soluble fraction of the SLG has been analyzed by mass spectrometry and was identified as archaetidic acid (25). However, as the lipid has been released from the protein by alkaline hydrolysis, this procedure may have hydrolyzed and thus removed the ethanolamine headgroup.

While investigating the interdependence between ArtA and HvPssA/HvPssD, we observed the recruitment of these proteins to the mid cell in *H. volcanii* (Fig. 4A). Considering these data and the observed mid-cell localization of the SLG (Fig. 4C), we propose a model for S-layer assembly, lipidation and growth in haloarchaea (Fig. 5B). First, SLG is recruited to mid cell, where it is transported across the cytoplasmic membrane in a Sec-dependent manner (26). Following secretion, SLG is processed and linked to archaetidylethanolamine by ArtA, requiring HvPssA/HvPssD for archaetidylethanolamine synthesis and/or interaction with ArtA.

There are still key aspects of haloarchaeal growth and shape control that are not addressed by our model. For example, it is still not clear how the deletion of either *artA*, *hvpssA*, or *hvpssD* generates a rod-shaped cell population (Fig. 2C). Furthermore, although it has been shown that the S-layer is not essential in other archaea (27, 28), overexpression of our SLG-msfGFP fusion drastically impacted the morphology of *H. volcanii* cells (Fig. 4D) beyond the lack of SLG processing and lipidation (11).

Therefore, it is possible that our SLG-msfGFP fusion is actually blocking the transport of or interaction with other yet unknown surface proteins essential for shape maintenance. The concept of having different classes of surface-modifying proteins counteracting each other has been demonstrated in bacteria, where different classes of Penicillin Binding Proteins (PBPs) act on the peptidoglycan cell wall to control cell width homeostasis in rod-shaped cells (29).

Interestingly, just like ovococcoid bacteria that are capable of mid-cell elongation and also lack a clear dedicated elongation machinery, haloarchaea may be also employing cytoskeletal polymers to direct different sub-complexes for cell elongation and cell division (30). This evidence is even more striking for coccoid bacteria, for which a single point mutation in FtsZ is able to induce cell elongation in *Staphylococcus aureus* (31). However, haloarchaea might have conserved at least two distinct elongation modes in addition to cell division, generating disk-like and rod-like populations (32). This scenario would also corroborate the morphological malleability of haloarchaea, being capable of assuming unusual shapes like triangles and squares (33, 34).

In spite of the bacteria that use the localization of specialized proteins to promote mid-cell elongation, it is important to point out that these mechanisms are likely not evolutionary related to the proposed haloarchaeal S-layer lipidation and cell elongation. First, even though there are examples of bacterial species with S-layer that carry out peptidoglycan cell-wall synthesis at mid cell (6), their new S-layer material is inserted as patches distributed all around the cell (35, 36). Second, lack of conservation in the protein architecture between archaeal and bacterial S-layers argue that they may have emerged independently of each other (8, 37).

In conclusion, by applying a set of different experimental approaches, we have confirmed that two putative lipid biosynthesis enzymes, HvPssA and HvPssD, are involved in the proteolytic cleavage and lipid labeling of ArtA substrates specifically at mid cell. We have also proposed, to the best of our knowledge, the first molecular model for archaeal cell elongation.

## Materials and Methods

### Strains and growth conditions

The plasmids and strains used in this study are listed in Table S1. *H. volcanii* strain H53 and its derivatives were grown at 45°C in semi-defined casamino acid (CA) medium supplemented with tryptophan (50 µg ml^−1^ final concentration) (38). Cells were cultivated either in liquid medium (orbital shaker at 250 rpm) or on solid 1.5% agar. Difco agar and Bacto yeast extract were purchased from Becton, Dickinson, and Company. Peptone was purchased from Oxoid. To ensure equal agar concentrations in all plates, agar was completely dissolved in the media prior to autoclaving, and autoclaved media were stirred before plates were poured. *Escherichia coli* strains were grown at 37°C in NZCYM medium (Fisher Scientific) supplemented with ampicillin (100 µg/ml).

### Plasmid preparation and *H. volcanii* transformation

DNA polymerase, DNA ligase, and restriction enzymes were purchased from New England BioLabs. Plasmids were initially transformed into *E. coli* DH5α cells. Plasmid preparations were performed using the QIAprep^®^ Spin Miniprep (Qiagen) kits. Prior to *H. volcanii* transformation, plasmids were transformed into the *dam*^−^ *E. coli* strain DL739. *H. volcanii* transformations were performed using the polyethylene glycol (PEG) method (38). All primers used to construct the recombinant plasmids are listed in Table S2.

### Generation of chromosomal *hvpssA* and *hvpssD* deletions in H53

Chromosomal deletions were generated by homologous recombination (pop-in pop-out) as previously described (19). Plasmid constructs for use in the pop-in pop-out knockout process were generated by using overlap PCR as described previously (39) as it follows: approximately 700 nucleotides flanking the *hvpssA* gene were PCR amplified and cloned into the haloarchaeal suicide vector pTA131. The *hvpssA* upstream flanking region is amplified with primers FW_pssA_KO_XbaI and RV_pssA_up while the *hvpssA* downstream flanking region is amplified using FW_pssA_dw and RV_pssA_KO_XhoI (primers are listed in Table S2). The *hvpssA* upstream and downstream flanking DNA fragments were fused by PCR using primers FW_pssA_KO_XbaI and RV_pssA_KO_XhoI, followed by cloning into pTA131 digested with *Xba*I and *Xho*I. The insertion of the correct DNA fragment into the cloning site of the recombinant plasmid was verified by sequencing using the same primers. The final plasmid construct, pFH38, contained upstream and downstream *hvpssA* flanking regions and was transformed into the parental H53 *H. volcanii* strain. To confirm the chromosomal replacement event at the proper location on the chromosome, colonies derived from these techniques were screened by PCR using the FW_pssA_KO_XbaI and RV_pssA_KO_XhoI primers. The *hvpssA* deletion mutant generated in strain H53 was designated FH38 (Table S1). For the generation of plasmid construct for chromosomal *hvpssD* deletion, approximately 700 nucleotides flanking the *hvpssD* gene were PCR amplified and cloned into the haloarchaeal suicide vector pTA131. The upstream flanking region was amplified with primers FW_pssD_KO_XbaI and RV_pssD_up while the downstream flanking region was amplified using FW_pssD_dw and RV_pssD_KO_XhoI. The flanking DNA fragments were fused by PCR using primers FW_pssD_KO_XbaI and RV_pssD_KO_XhoI, followed by cloning into pTA131 digested with *Xba*I and *Xho*I. The insertion of the correct DNA fragment into the cloning site of the recombinant plasmid was verified by sequencing using the same primers. The final plasmid construct, pFH43, contained upstream and downstream *hvpssD* flanking regions and was transformed into the parental *H. volcanii* strain H53. Confirmation of *hvpssD* deletion on the chromosome was screened by PCR using the FW_pssD_KO_XbaI and RV_pssD_KO_XhoI primers. The *hvpssD* deletion mutant generated in strain H53 was designated FH63 (Table S1).

### Construction of expression plasmids for HvPssA and HvPssD

To construct a tryptophan-inducible *H. volcanii hvpssA* gene with C-terminal His-tag, its coding region was amplified by PCR using primers FW_pssA_OE_NdeI and RV_pssA_OE_EcoRI_His (Table S2). Meanwhile, for the construction of *H. volcanii hvpssD* gene with C-terminal His-tag, its coding region was amplified by PCR using primers FW_pssD_OE_NdeI and RV_pssD_OE_EcoRI_His (Table S2). The PCR product was cloned into the expression vector pTA963 that had been digested with *Nde*I and *Eco*RI. This places the *hvpssA* or *hvpssD* gene under the control of the inducible tryptophanase promoter (p.*tna*). The recombinant pTA963 carrying the *hvpssA* gene was designated pFH39 and the pTA963 carrying *hvpssD* was designated pFH44. To complement the Δ*hvpssA* strain FH38, this strain was transformed with plasmid pFH39 to result in FH56. For complementation of Δ*hvpssD* strain FH63, this strain was transformed with plasmid pFH44 to result in FH70. The H53, Δ*hvpssA*, and Δ*hvpssD* strains were also transformed with the empty expression vector pTA963 which was used as a control.

### Construction of expression plasmid for *csg*-msfGFP(SW) and msfGFP

To construct a tryptophan-inducible *H. volcanii csg* tagged with msfGFP, a sandwich fusion was created by intercalating msfGFP gene product amplified from synthetic fragment (oligos oHV81 and oHV82) in between the first 102bp (the secretion signal sequence of SLG, oligos oAB500 and oAB501) and the rest of the *csg* coding region (oligos oHV83 and oAB502). The 3 fragments were then assembled by Gibson assembly (40) together with the pTA962 plasmid (41), digested with NheI and BamHI and transformed into DH5α cells. The same process was followed for the cloning of msfGFP alone (oligos oHV101 and oHM68). The clones were subjected to validation by PCR and Sanger sequencing (Table S1).

### Generation of chromosomal msGFP fusions in H26

C-terminal translational fusion constructs, including a pyrE2 cassette for selection, were generated by direct transformation of PCR fragment ensembles generated by Gibson assembly (40), transformed directly in *H. volcanii* H26 cells and selected by growth in the absence of uracil. PCR fragments from ftsZ*1* (oligos oHV3 and oHV4 for upstream region and oHV8 and oHV9 for downstream region), *artA* (oligos oHM91 and oHM92 for upstream region and oHM93 and oHM94 for downstream region), *hvpssA* (oligos oHV156 and oHV157 for upstream region and oHV158 and oHV159 for downstream region), and *hvpssD* (oligos oHV126 and oHV127 for upstream region and oHV128 and oHV129 for downstream region) were assembled to msfGFP (oligos oHM34 and oHM6) and the pyrE2 cassette (oligos oHV6 and oHV7). Chromosomal replacements were confirmed by PCR and Sanger sequencing (Table S1).

### Immunoblotting

Liquid cultures were grown until mid-log phase (OD_600_ 0.2-0.5) and the cells were harvested by centrifugation at 3,800 × g for 5 min at room temperature. Cell pellets were resuspended and lysed in 1% (v/v) NuPAGE lithium dodecyl sulfate (LDS) sample buffer supplemented with 100 mM dithiothreitol (DTT) and stored at −20°C. Samples were electrophoresed on 4-12% Bis-Tris polyacrylamide gels (Invitrogen) with NuPAGE 3-(N-morpholino) propane sulfonic acid (MOPS) sodium dodecyl sulfate (SDS) running buffer (Invitrogen). Proteins were then transferred to polyvinylidene difluoride (PVDF) membranes (Millipore) using a semi-dry transfer apparatus at 15 V for 30 minutes (Bio-Rad). Subsequently, the membrane was washed twice in PBS, blocked for one hour in 3% bovine serum albumin (BSA) in PBS, and washed twice in PBS with 1% Tween-20 and once with PBS. For detection of the poly His-tag, the mouse anti-penta-His antibody (Qiagen; Catalog #34660) was used at a 1:2,000 dilution in 3% BSA in PBS with sodium azide. For the secondary antibody, HRP-conjugated Amersham ECL anti-mouse IgG from sheep (GE) was used at a 1:20,000 dilution in 10% nonfat milk in PBS. For detection of HVO_0405_Nterm (see Suppl Text S1 and associated methods), the rabbit anti-HVO_0405-N-term serum (12) was used as the primary antibody at a 1:10,000 dilution in 3% BSA in PBS with sodium azide. For the secondary antibody, HRP-conjugated Amersham ECL anti-rabbit IgG from donkey (GE) was used at a 1:60,000 dilution in 10% nonfat milk in PBS.

### Lipid radiolabeling

The H53 parent strain carrying the vector control pTA963, and the Δ*hvpssA* deletion strain carrying either the *hvpssA* expression plasmid pFH39 or the vector control pTA963 were grown in 5 ml liquid CA medium. Upon reaching mid-log phase (OD_600_ ~0.5), 20 µl of each culture was transferred into 1 mL of fresh liquid CA medium supplemented with [^14^C] mevalonic acid (resuspended in ethanol) at a final concentration of 1 µCi/ml. The cultures were harvested after reaching mid-log phase and proteins were precipitated from 1 ml cultures with 10% TCA, followed by a delipidation step to remove non-covalently linked lipid as described previously (42, 43). The delipidated proteins were separated by 7% Tris-Acetate (TA) LDS-PAGE. For analysis of the samples, the gel was dried onto blotting paper using a Gel Dryer (Bio-Rad model 583), exposed to a phosphor screen (Molecular Dynamics) for 3 weeks, and analyzed using a Typhoon imager (Amersham Biosciences).

### Motility assays

The motility assays of *H. volcanii* H53 (parent), Δ*artA*, Δ*hvpssA*, and Δ*hvpssD* strains carrying the plasmid expressing the complementary gene (or pTA963 as control) were performed on 0.35% agar in CA medium supplemented with tryptophan as described previously (39). A toothpick was used to stab inoculate the agar, followed by incubation at 45°C. Halo sizes around the stab-inoculation site were measured after 3–5 days of incubation.

### Growth curves

Growth curves were measured using a Biotek PowerWaveX2 microplate spectrophotometer. *H. volcanii* H53 (parent), Δ*artA*, Δ*hvpssA*, and Δ*hvpssD* strains carrying the plasmid expressing the complementary gene (or pTA963 as control) were first incubated in 5 ml liquid cultures in CA medium supplemented with tryptophan with continuous shaking at 45°C, until suitable OD_600_ values (0.2–0.5) were reached. Approximately 6 μl of each culture (adjusted to correct for OD_600_ differences) were then transferred into 194 μl of fresh CA medium supplemented with tryptophan (50 µg ml^−1^ final concentration) and grown to stationary phase, with OD_600_ recordings taken every 30 min.

### Light microscopy

The *H. volcanii* strains H53 (parent), Δ*artA*, Δ*hvpssA*, and Δ*hvpssD* carrying the plasmid expressing the complementary gene (or pTA963 as control) were inoculated from colony to 5 ml CA liquid medium and grown until they reached mid-log phase (OD_600_ ~0.4-0.5). Serial liquid to liquid sub-inoculations were carried out by transferring 10 µl of the liquid culture to 5 ml fresh liquid CA medium with up to two transfers. Subsequently, 1 ml of each culture was concentrated by centrifugation at 4,911 × g for 1 min, pellets resuspended in 10 µl of liquid CA medium. Then, 10 µl of the concentrated cells was transferred to under a 0.5% agarose pad with CA medium and observed using a Nikon Eclipse TiE inverted TIRF microscope. ArtA-msfGFP time lapses were acquired by culturing *H. volcanii* cells inside Millipore ONIX CellAsic microfluidic plates as previously described (33). Images were taken under 5-minute intervals for 12 hours in both phase contrast and with 488 nm laser channels. *H. volcanii* protoplasts were generated within microfluidic channels by the addition of 1 mg/mL of Proteinase K (Invitrogen) in YPC medium until cells lost shape. Subsequently, the cells were washed with fresh YPC and time lapse was recorded with 10-minute intervals for 12 hours.

## Supporting information

Abdul-Halim 2019_Supplementary Data

Abdul-Halim 2019_Movie S1

Abdul-Halim 2019_Movie S2

## Acknowledgments

We want to thank Howard Goldfine, the Pohlschroder, Daldal and Garner labs for helpful discussions. We also want to thank Henry Miziorko, Ethan Garner and Jenny Zheng for access to equipment and reagents. MP and MAH were supported by the National Science Foundation grant 1817518. SS was supported by the German Science Foundation Postdoctoral Fellowship.

