## Supplementary material for "LIPID ANCHORING OF ARCHAEOSORTASE SUBSTRATES AND MID-CELL GROWTH IN HALOARCHAEA": Abdul-Halim 2019_Supplementary Data

**This document includes:**

Figures S1 to S3
Tables S1 and S2
Movie captions S1 and S2

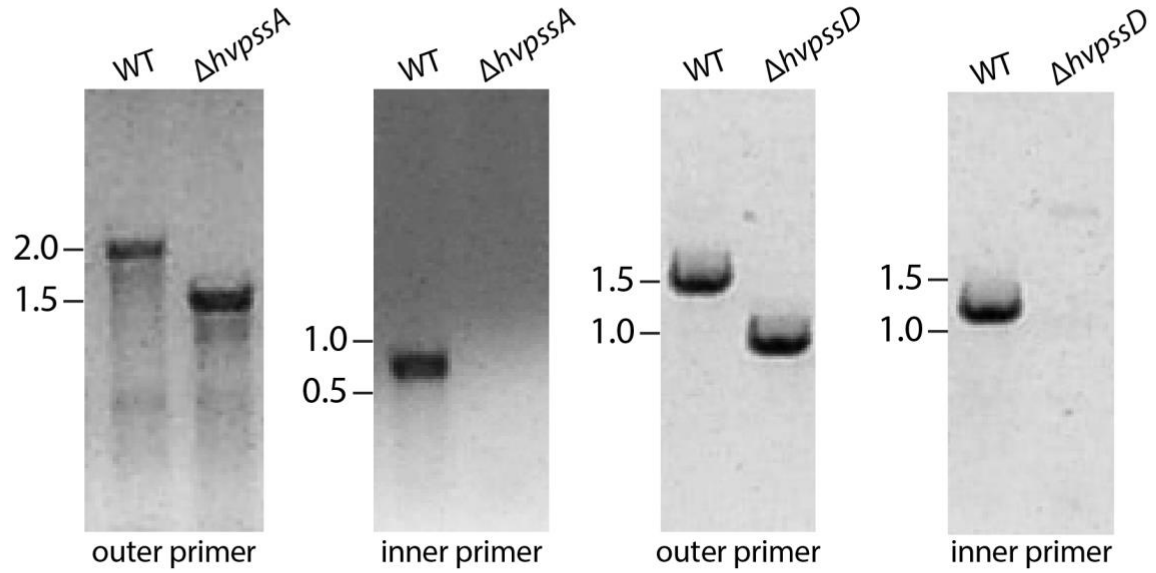


**Figure S1: *H. volcanii* HvPssA and HvPssD are not essential under standard laboratory growth conditions.** PCR amplification using primers against the flanking regions located approximately 700 bp upstream and 700 bp downstream of *hvpssA* or *hvpssD* (outer primer) and primers specific for the *hvpssA* or *hvpssD* encoding gene (inner primer). The template DNA was isolated from H53 wild-type (WT), Δ*hvpssA*, and Δ*hvpssD* strains.

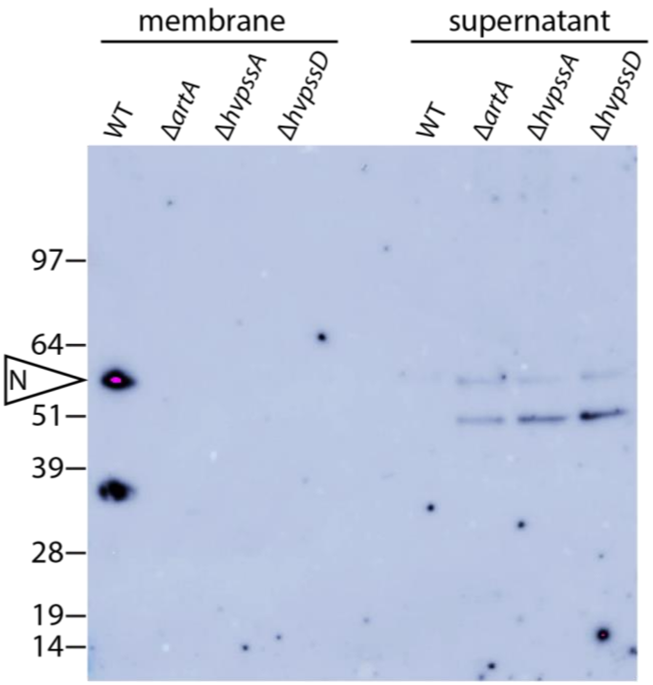


**Figure S2: Lack of lipidation prevents ArtA-dependent HVO_0405 processing** . Western blot analysis of membrane (m) and supernatant (s) fractions of H53 (wt), Δ*artA*, Δ*hvpssA*, and Δ*hvpssD* strains expressing, *in trans*, HVO_0405-6xHis. The N-terminal domain of HVO_0405 was detected using anti-HVO_0405-N-term antibodies. The processed N-terminal HVO_0405, marked as "N", can only be detected in the membrane fraction of the wt. Bands in the supernatant fraction of all mutant strains are not related to ArtA-dependent processing (i.e. cleaved but not lipidated HVO_0405), as they are detected for Δ*artA* as well. Instead, they probably indicate proteolytic degradation products of HVO_0405. Numbers indicate molecular mass in kilodaltons.

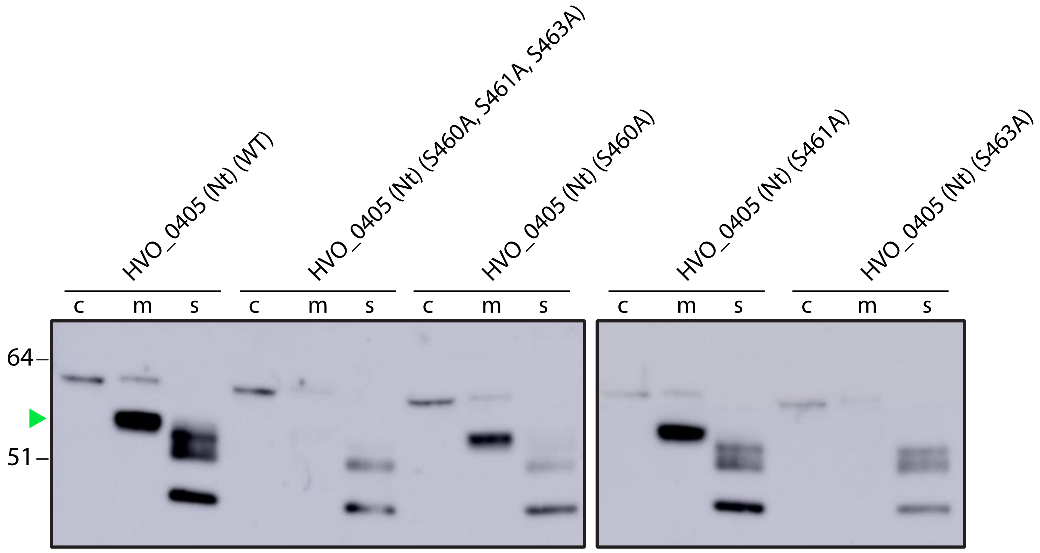


**Figure S3: HVO_0405 Ser$_{463}$ residue is critical for the substrate C-terminal processing.** Western blot analysis of protein from cytoplasmic (c), membrane (m), and supernatant (s) fractions of the Δ*hvo_0405* mutant transformed with pTA963 expressing the His-tagged N-terminal domain of HVO_0405 ("HVO_0405(N)") or mutants thereof. The HVO_0405 N-terminal domain reverts the strain-specific domain fusion event. This domain is either wild-type HVO_0405(N)-6XHis, or carries point mutation of selected Ser residues to Ala. We generated a triple mutant (Ser$_{460}$ Ser$_{461}$ Ser$_{463}$) and single mutants for each of these Ser residues. Anti-HVO_0405-N-term antibodies (12) were used to determine the HVO_0405 C-terminal processing. Faster migrating C-terminally processed mature HVO_0405 are indicated with a green arrowhead. Numbers indicate molecular masses in kDa. The images shown represent the results from at least three independent replicates.

First frame snapshot

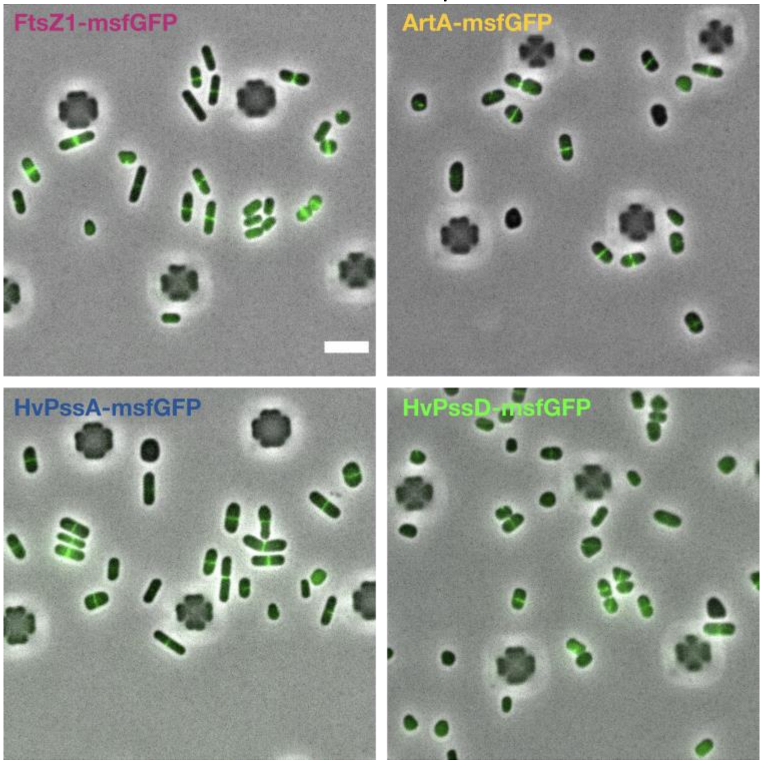


**Movie S1: Movie S1: FtsZ1, ArtA, HvPssA and HvPssD mid-cell localization and dynamics throughout the cell cycle in *H. volcanii*.** Time lapses were acquired with cells growing inside CellAsic microfluidic device. Images of merged phase contrast (grey) and FITC (green) were taken every 5 minutes for 12 hours. Scale represents 5µm.

First frame snapshot

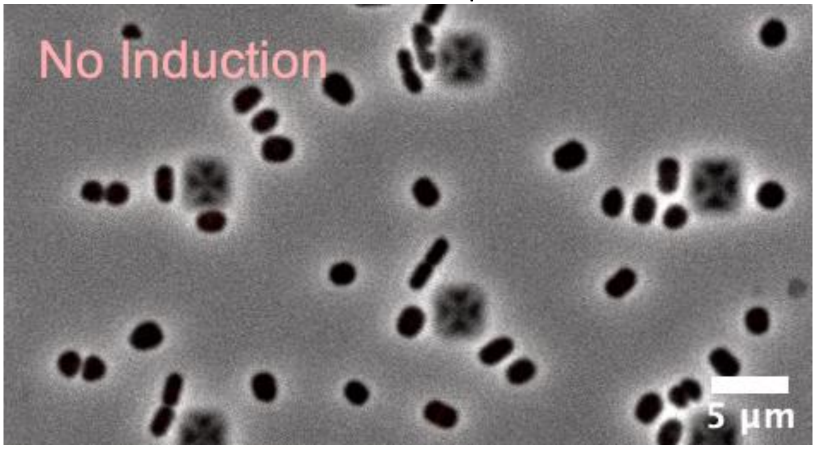


**Movie S2: SLG-msfGFP mid-cell localization in *H. volcanii* cells.** Time lapses were acquired with cells growing inside CellAsic microfluidic device. Images of merged phase contrast (grey) and FITC (red) were taken every 5 minutes for 12 hours. During the first 6 hours, cells were grown and imaged in absence of Tryptophan (No Induction). Between 6 and 12 hours, it was added Tryptophan to a final concentration of 500 mM, undicing the SLG-msfGFP expression (Induction). Scale bar represents 5 µm.

**Table S1: Strains and plasmids used in this work**

| Alias | Genotype/Description | Source |
|---|---|---|
| **Plasmids** | | |
| pTA131 | $Amp_r$; pBluescript II with BamHI-XbaI fragments from pGB70 harboring *p.fdx-pyrE2* | (19) |
| pTA963 | $Amp_r$, *pyrE2* and *hdrB* markers, Trp-inducible (*p.tna*) promoter | (41) |
| pFH25 | pTA963 carrying *hvo_0405* with C-terminal 6×His-tag | (12) |
| pFH38 | pTA131 carrying 700 bp upstream and 700 bp downstream *hvpssA* flanking region | This work |
| pFH43 | pTA131 carrying 700 bp upstream and 700 bp downstream *hvpssD* flanking region | This work |
| pFH39 | pTA963 carrying *hvpssA* with C-terminal 6×His-tag | This work |
| pFH44 | pTA963 carrying *hvpssD* with C-terminal 6×His-tag | This work |
| pFH51 | pTA963 carrying *hvo_0405* N-terminal LVIVD-domain with C-terminal 6×His-tag with $Ser_{460}$, $Ser_{461}$, and $Ser_{463}$ mutated to Alanine | This work |
| pFH52 | pTA963 carrying *hvo_0405* N-terminal LVIVD-domain with C-terminal 6×His-tag with $Ser_{460}$ mutated to Alanine | This work |
| pFH53 | pTA963 carrying *hvo_0405* N-terminal LVIVD-domain with C-terminal 6×His-tag with $Ser_{461}$ mutated to Alanine | This work |
| pFH54 | pTA963 carrying *hvo_0405* N-terminal LVIVD-domain with C-terminal 6×His-tag with $Ser_{463}$ mutated to Alanine | This work |
| pFH55 | pTA963 carrying *hvo_0405* N-terminal LVIVD-domain with C-terminal 6×His-tag | This work |
| **Strains** | | |
| DH5α | *E. coli* F− ϕ80dlacZΔM15 (*lacZYA-argF*) U169 *recA1 endA1 hsdR17*(*rK− mK−*) *phoA supE44 thi-1 gyrA96 relA1* | Invitrogen |
| H26 | *H. volcanii* Δ*pyrE2* | (19) |
| H53 | *H. volcanii* Δ*pyrE2* Δ*trpA* | (19) |
| FH27 | H53<br>pTA963::*hvo_0405*-6×His | (12) |
| FH28 | H53 Δ*artA*<br>pTA963::*hvo_0405*-6×His | (12) |
| FH55 | H53 Δ*hvpssA*<br>pTA963 | This work |
| FH56 | H53 Δ*hvpssA*<br>pTA963::*hvpssA*-6×His | This work |
| FH57 | H53 Δ*hvpssA* | This work |

| | | |
|---|---|---|
| | pTA963::*hvo_0405*-6×His | |
| FH69 | H53 Δ*hvpssD*<br>pTA963 | This work |
| FH70 | H53 Δ*hvpssD*<br>pTA963::*hvpssD*-6×His | This work |
| FH71 | H53 Δ*hvpssD*<br>pTA963::*hvo_0405*-6×His | This work |
| FH73 | H53 Δ*hvo_0405*<br>pTA963::N-term-*hvo_0405*$_{S460A,S461A,S463A}$-6×His | This work |
| FH74 | H53 Δ*hvo_0405*<br>pTA963::N-term-*hvo_0405*$_{S460A}$-6×His | This work |
| FH75 | H53 Δ*hvo_0405*<br>pTA963::N-term-*hvo_0405*$_{S461A}$-6×His | This work |
| FH76 | H53 Δ*hvo_0405*<br>pTA963::N-term-*hvo_0405*$_{S463A}$-6×His | This work |
| FH77 | H53 Δ*hvo_0405*<br>pTA963::N-term-*hvo_0405*-6×His | This work |
| aBL128 | Δ*pyrE2*<br>*artA::artA-msfGFP-pyrE2* | This work |
| aBL129 | Δ*pyrE2*<br>*hvpssD::hvpssD-msfGFP-pyrE2* | This work |
| aBL183 | Δ*pyrE2*<br>*hvpssA::hvpssA-msfGFP-pyrE2* | This work |
| aBL131 | Δ*pyrE2*<br>*ftsZ1::ftsZ1-msfGFP-pyrE2* | This work |
| aBL184 | Δ*pyrE2*<br>pTA962::msfGFP(SW) | This work |
| aBL118 | Δ*pyrE2*<br>pTA962::*csg-msfGFP*(SW) | This work |

**Table S2: Oligos used in this work**

| Alias | 5’→3’ Sequence | Source |
|---|---|---|
| FW_pssA_OE_NdeI (IP) | ATTAATCATATGATGAGACCCCGATTCG | This work |
| RV_pssA_OE_EcoRI_His (IP) | TATATTGAATTCTCAGTGATGGTGATGGTGATGGTG ATGGTGATGCGGGCCGCCCGCCCAGTAGAAGCG | This work |
| FW_pssA_KO_XbaI (OP) | AATAAAATCTAGAGGCTTTTTATGCGACCGCGCTTC GAGC | This work |
| RV_pssA_KO_XhoI (OP) | AATAAAACTCGAGCCGCGAGCCACGCGTGGTACG | This work |
| RV_pssA_up | CCGTTCCGCCGCCCCGGACCGATGCGATTGCGTACC TGCCC | This work |
| FW_pssA_dw | GGGCAGGTACGCAATCGCATCGGTCCGGGGCGGCG GAACGG | This work |
| FW_pssD_KO_XbaI (OP) | TATATTTCTAGAGTCCGCCGGCTACGC | This work |
| RV_pssD_KO_XhoI (OP) | TATATTCTCGAGCCGTACAGGTCGGCGTAG | This work |
| RV_pssD_up | AGACGGCCAACGCCGACAGCGACGCTTATGCCTCGC CGCGGACGTC | This work |
| FW_pssD_dw | GACGTCCGCGGCGAGGATAAGCGTCGCTGTCGGCGT TGGCCGTCT | This work |
| FW_pssD_OE_NdeI (IP) | TATATTCATATGATGCGGTTCGCACCC | This work |
| RV_pssD_OE_EcoRI_His (IP) | TATATTGAATTCTCAGTGATGGTGATGGTGATGCTCC CGCCGCGCCAA | This work |
| FW_0405_NdeI | TATATTCATATGATGGACCGCCGCCAGTT | This work |
| RV_0405_S460,S461,S463 A_upstream | CGTCGCGGTCGCCGCTTCGGTTTCGGACGCCGA | This work |
| RV_0405_S460A_upstrea m | CGTCGAGGTCGACGCTTCGGTTTCGGACGCCGA | This work |
| RV_0405_S461A_upstrea m | CGTCGAGGTCGCCGATTCGGTTTCGGACGCCGA | This work |
| RV_0405_S463A_upstrea m | CGTCGCGGTCGACGATTCGGTTTCGGACGCCGA | This work |
| FW_0405_S460,S461,S463 A_downstream | GAAGCGGCGACCGCGACGACCGACGCGCCGG | This work |
| FW_0405_S460A_downstr eam | GAAGCGTCGACCTCGACGACCGACGCGCCGG | This work |

| | | |
|---|---|---|
| FW_0405_S461A_downstream | GAATCGGCGACCTCGACGACCGACGCGCCGG | This work |
| FW_0405_S463A_downstream | GAATCGTCGACCGCGACGACCGACGCGCCGG | This work |
| RV_0405_LVIVD_His_EcoRI | TATATTGAATTCTCAGTGATGGTGATGGTGATGCGGGCCGCCACTCCGATGCTTCGA | This work |
| pTA963-1F | CACACACCAGTCCACGAG | This work |
| pTA963-1R | CGCAATTAACCCTCACTAAAG | This work |
| oHV3 | CGTCCTCCGTAAACCG | This work |
| oHV4 | GTCCGCTACCCTCAAGCTCGACGTAGTCGATGTCT | This work |
| oHV6 | TTAGCCGTCGGCGTC | This work |
| oHV7 | CGTGGATAAAACCCCTCG | This work |
| oHV8 | CGAGGGGTTTTATCCACGTCGAGCCGTCCC | This work |
| oHV9 | CGAAGAAACGGTTTTGTGG | This work |
| oHV81 | CTCGAGGGATCTGGC | This work |
| oHV82 | CCCACTGCCTTGACC | This work |
| oHV83 | GGTCAAGGCAGTGGGGAGCGTGGAAACCTCG | This work |
| oHV101 | CGGACCTATTGCGCATATGCGAAAAGGGGAAGAATTGTTTAC | This work |
| oHV126 | GCTCAAGGAGTCCGC | This work |
| oHV127 | GTCCGCTACCCTCAAGCTCCCGCCGCG | This work |
| oHV128 | CGAGGGGTTTTATCCACGGCGTCGCTGTCGG | This work |
| oHV129 | GCAGGTGTCGATTGCC | This work |
| oHV156 | CGACGACGATGGTGC | This work |
| oHV157 | GTCCGCTACCCTCAAGCGCCCAGTAGAAGCG | This work |
| oHV158 | CGAGGGGTTTTATCCACGCCGGGGCGGC | This work |
| oHV159 | CCGAACTCCGCTTCG | This work |
| oHM6 | CGCCGACGGCTAATCATTTGTAAAGTTCATCCATTCCATGC | This work |
| oHM34 | CTTGAGGGTAGCGGAC | This work |
| oHM68 | GGTGCGGCCGCTCATTTGTAAAGTTCATCCATTCCA | This work |

| | | |
|---|---|---|
| oHM91 | AGTACGTATGCCCGGT | This work |
| oHM92 | CGAGTTAGGGCTCGACCTTGAGGGTAGCGGAC | This work |
| oHM93 | CGAGGGGTTTTATCCACGTTACGCCGTCGGAGT | This work |
| oHM94 | CCACATGTTCAGCATATCGG | This work |
| oAB500 | CGGACCTATTGCGCATATGACAAAGCTCAA | This work |
| oAB501 | GCCAGATCCCTCGAGCGCGGCGGCACTTCC | This work |
| oAB502 | CAGAGGTGCGGCCGCTTAGTTCTCGCGGCG | This work |
